# The Prion-Like Properties of the INO80 Chromatin Remodeler Regulate Metabolic Gene Expression

**DOI:** 10.64898/2026.05.18.722789

**Authors:** Dylan C. Englund, Keith C. Garcia, Daniel F. Jarosz, Ashby J. Morrison

## Abstract

The INO80 chromatin-remodeling complex is a multi-subunit regulator of DNA-templated processes, yet the mechanisms that control remodeler function *in vivo* are not completely known. In this study, we report that the Ies6 subunit of the INO80 complex encodes a prion-like domain (PLD) within a larger region of predicted disorder. After transient inducible overexpression, both full-length Ies6 and the PLD domain alone aggregate in a manner that evades proteasome-mediated degradation, suggestive of prion-like behavior. Cytosolic aggregates are also visible with fluorescent microscopy following expression of the PLD alone. Loss of the protein chaperone Hsp70 increases Ies6 aggregation. In addition, deletion of *ARP5*, another INO80 subunit and binding partner of Ies6, also results in elevated protein aggregation. Finally, transcriptome analysis indicates that loss of PLD-mediated aggregation alters the expression of telomere-proximal and metabolic stress-responsive genes, including those involved in glucose starvation and respiratory programs, even under glucose-replete conditions. Together, these findings support a model in which Ies6 couples the INO80 complex to metabolic gene regulation via prion-like behavior, providing a potential new mechanism for tuning chromatin-based metabolic adaptation.

## INTRODUCTION

Eukaryotic transcriptional programs depend on the ability of chromatin to reorganize rapidly in response to environmental and developmental cues. Chromatin remodeling complexes provide one major mechanism for this regulation by using the energy of ATP hydrolysis to reposition, evict, or exchange nucleosomes, thereby altering accessibility of DNA to transcriptional regulators. For example, the INO80 complex is an evolutionarily conserved remodeler that organizes promoter-proximal nucleosome architecture (Bowman 2025). Furthermore, INO80 regulates gene expression in both metabolic pathways in yeast and developmental pathways in mice (Yao et al. 2016; Gowans et al. 2018; Rhee et al. 2018, 2021). Thus, the INO80 complex is an important regulator of inducible transcriptional programs that are critical for metabolic homeostasis and proper developmental timing.

Both the mammalian and *S. cerevisiae* INO80 complexes are large megadalton multi-subunit assemblies organized around the Snf2-family ATPase Ino80, which serves as a central scaffold for several functionally distinct subunit modules (Shen et al. 2000; Tosi et al. 2013; Chen et al. 2011). Interestingly, in *S. cerevisiae*, the Arp5 (actin-related protein 5) and Ies6 (Ino-eighty subunit 6) subunits have been found to form an abundant subcomplex *in vivo* that can dynamically associate with the larger INO80 complex *in vitro* to stimulate ATP hydrolysis and nucleosome sliding that are central to transcriptional processes (Yao et al. 2016, 2015).

Transcriptional profiling of *arp5Δ* and *ies6Δ* mutants exhibit de-repression of oxidative phosphorylation genes, leading to constitutively elevated mitochondrial potential and oxygen consumption (Yao et al. 2016, 2015). An expansive genetic interaction screen further identified the Ies6 module as a hub linking INO80 to metabolic homeostasis and showed that INO80 relays TORC1 signaling to chromatin by sustaining histone acetylation at TOR-responsive genes (Beckwith et al. 2018). Consistent with this, INO80 is required for the periodic oscillations of respiration and gene expression that define the Yeast Metabolic Cycle, and its loss uncouples cell division from nutrient availability (Gowans et al. 2018). Collectively, these studies position the INO80 complex, and the Arp5–Ies6 module in particular, as a key chromatin-based integrator of metabolic state and gene expression.

Another mechanism by which cells coordinate adaptive responses in changing metabolic environments is through prion-like conformational switching. Although such conformational switches were initially regarded as pathological, current evidence supports adaptive roles for prion-like assemblies in order to access alternative phenotypic states during environmental stress (Wickner 1994; Alberti et al. 2009). For example, the [*GAR+*] prion relaxes glucose repression, allowing yeast and other fungi to use alternative carbon sources in the presence of glucose and thereby shift from metabolic specialists toward metabolic generalists (Brown and Lindquist 2009; Jarosz et al. 2014; Garcia et al. 2016). Like other prions, [*GAR+*] assembly and inheritance are regulated by chaperones, specifically heat shock proteins, which mediate protein conformational change in order to provide a rapid and reversible route for adapting cellular physiology to changing nutrient environments (Brown and Lindquist 2009; Shorter and Lindquist 2005; Chernoff et al. 1995).

Transcriptional changes driven by prions can be linked to chromatin modification. For example Snt1, the scaffold subunit of the Set3C histone deacetylase complex can form a prion state, termed [*ESI+*] for “expressed sub-telomeric information” (Harvey et al. 2020). [*ESI+*] establishes transgenerational inheritance of an active chromatin state across normally repressed subtelomeric regions, resulting in gene activation and stress resistance. Additionally, Swi1, a subunit of the SWI/SNF chromatin remodeling complex, forms the [*SWI+*] prion, producing heritable changes in SWI/SNF-dependent gene regulation (Du et al. 2020, 2008). Together, [*ESI+*] and [*SWI+*] demonstrate that yeast prion-like domains can create reversible regulatory states that connect environmental stress and transcriptional control through chromatin-based adaptation.

In this study, we investigate the INO80 subunit Ies6 and find that it exhibits prion-like properties, namely protein aggregation that is regulated by the Hsp70 chaperone and the INO80 subunit Arp5. Transcriptional assays reveal that loss of Ies6 aggregation alters gene expression in pathways related to energy metabolism. Collectively, these results describe a new prion-like mechanism of transcriptional regulation for the INO80 complex that is important for metabolic homeostasis.

## RESULTS

### The Ies6 subunit of the INO80 complex contains a predicted prion-like domain

Computational modeling identified an N-terminal region of 23 amino acids in Ies6 that is enriched in prion-forming amino acids, as well as a region of predicted prion-like propensity (Figure 1A). Both of these regions are within an amino acid region predicted to be disordered. This nested architecture, a compositionally biased prion-like sequence embedded in a longer disordered region, is characteristic of yeast prions and prion-like proteins, in which the disordered state of the prion-like domain enables conformational switching to self-templating aggregate assemblies (Alberti et al. 2009). (Note, the prion-like propensity prediction is likely an underestimate of this 23-amino acid domain, as it is calibrated on regions of more than 40 residues.)

**Figure 1.**
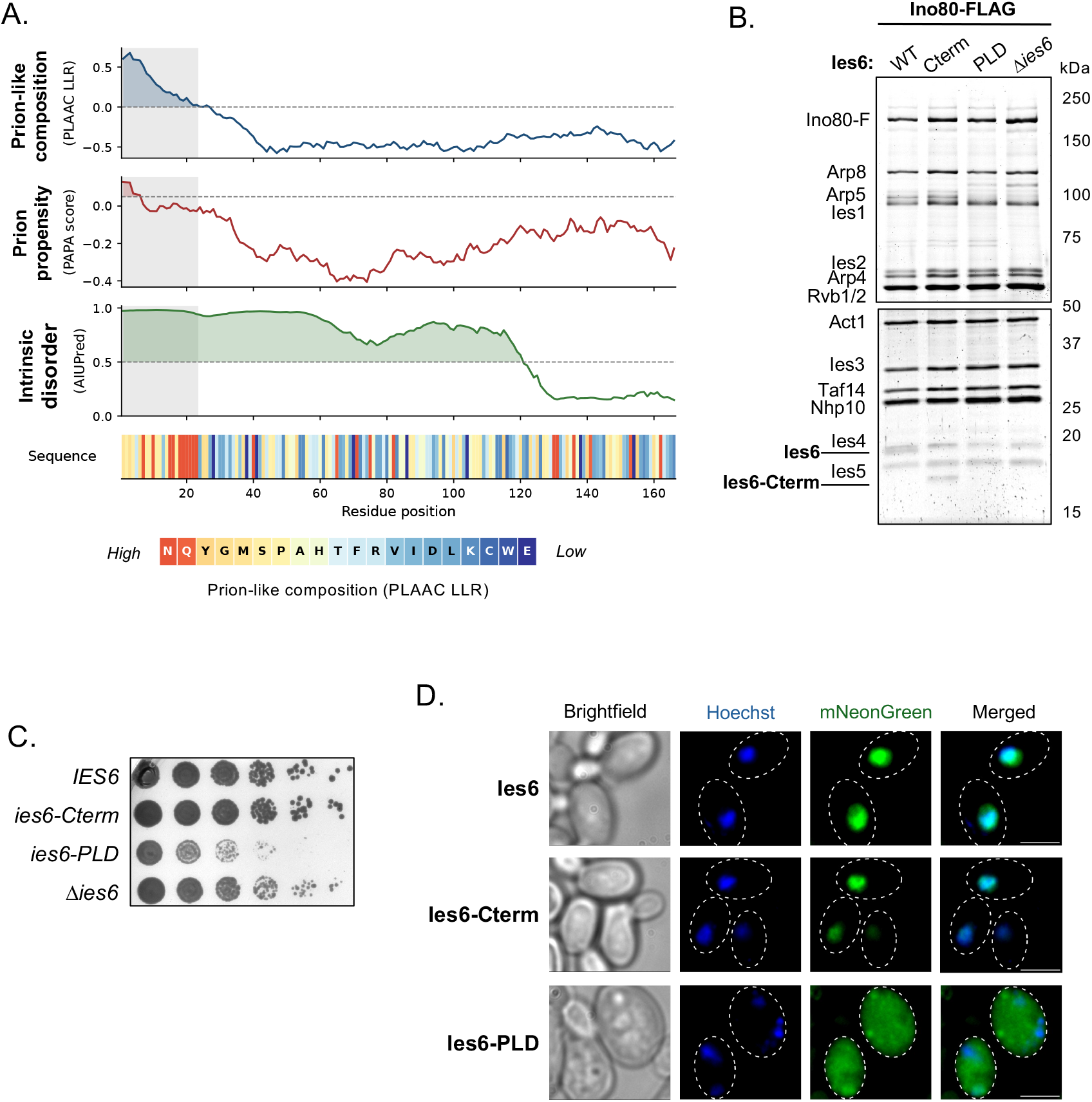
The Ies6 subunit of the INO80 complex contains a predicted N-terminus prion-like and disordered domain. **(A)** Ies6 per residue scores: prion-like amino acid composition (PLAAC log-likelihood ratio, LLR; blue) (Lancaster et al. 2014), prion propensity (PAPA score; red) (Toombs et al. 2010), and intrinsic disorder probability (AIUPred; green) (Erdős and Dosztányi 2024). Dashed horizontal lines indicate the standard significance thresholds. The grey vertical band denotes the prion-like domain (PrLD) boundaries assigned by the PLAAC hidden Markov model Viterbi parse (residues 1–23). The bottom sequence track shows each residue colored by its individual PLAAC LLR contribution. **(B)** Ino80-Flag purification with the indicated *ies6* mutants. Top, 6% SDS-PAGE gel and bottom, 13.5% SDS-PAGE gel. Previously identified INO80 subunits (Shen et al. 2000; Yao et al. 2015, 2016) are indicated on the side of the top 6% SDS-PAGE gel and the bottom 13.5% gel. **(C)** Fitness assays of indicated yeast strains grown on YPD. **(D)** Fluorescent imaging of indicated log-phase wild-type and mutant Ies6-mNeonGreen tagged strains with Hoechst DNA staining. Scale bar indicates 5 microns.

To identify features associated with the Ies6 prion-like domain (PLD), we generated two complementary truncation mutants: one containing only the predicted N-terminal prion-like domain amino acids 1-23 (*ies6-PLD*), and one mutant consisting of C-terminal amino acids 24-166 without the PLD domain (*ies6-Cterm*). Purification of INO80 complexes with these mutants demonstrates that the composition of the complex is unaltered in cells expressing Ies6 C-terminus (Figure 1B). However, when the Ies6-PLD alone is expressed, the association of Arp5 with the INO80 complex is greatly reduced. (Note, because of its small size, the PLD domain is not visible on standard glycine-based SDS-PAGE gels.) Similarly, complete deletion of *IES6* also results in the loss of Arp5 from the INO80 complex, consistent with previous results demonstrating that an intact Arp5-Ies6 subcomplex is needed for its association with the INO80 complex (Yao et al. 2015, 2016).

We then performed fitness assays and found that while *ies6-Cterm* cells exhibited growth similar to wildtype cells, the *ies6-PLD* cells exhibited viability defects that were more pronounced than that of *ies6Δ* cells (Figure 1C). Fluorescent imaging demonstrated that while wild-type Ies6 and the Ies6-Cterm mutant colocalize with DNA, the Ies6-PLD mutant is broadly distributed throughout the cell (Figure 1D). Together with the INO80 purifications, these results strongly suggest that the C-terminal domain is important for association with the INO80 complex and chromatin targeting, while the PLD may serve a regulatory role in Ies6 function.

### The prion-like domain of Ies6 promotes aggregation

To examine the behavior of Ies6’s PLD domain, we used a system that exploits the protease-resistant phenotypes previously observed for prion aggregates (Masison and Wickner 1995; McKinley et al. 1983). This system is composed of a fusion protein that combines Ies6 with a selection marker and a previously optimized destabilizing domain that targets proteins for proteasome-mediated degradation (Iwamoto et al. 2010; Yang et al. 2026) (Figure 2A). The fusion protein is under the control of a tightly-regulated Tet-inducible promoter for overexpression (Azizoglu et al. 2021). Cells are grown in the presence of doxycycline, followed by growth on selective media without doxycycline induction. Subsequent growth on selective media suggests propagation of prion-like aggregates that evade proteolytic clearance and confer selection resistance.

**Figure 2.**
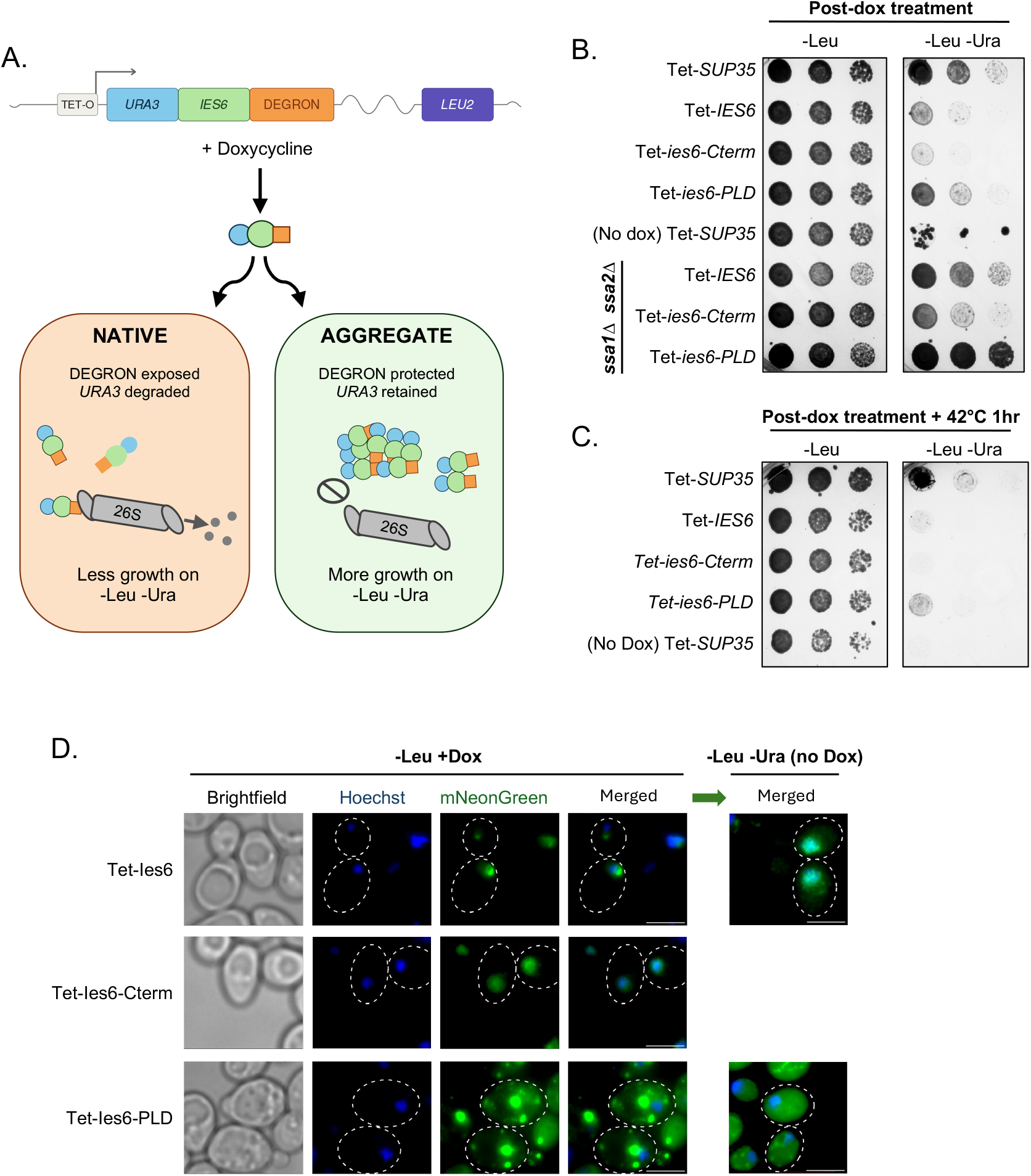
The prion-like domain (PLD) of Ies6 aggregates when overexpressed. **(A)** Illustration of Tet-inducible aggregation selection system. **(B)** Growth of wild-type and Hsp70 mutants (*ssa1Δ ssa2Δ*) expressing indicated Tet-inducible constructs on both -Leu (no aggregation selection) and -Leu -Ura (aggregation selection). All strains were treated with doxycycline prior to plating, except where indicated in the (no dox) negative control. **(C)** As in (B), but cells were heat shocked at 42°C for 1 hour prior to plating. **(D)** Fluorescent imaging of indicated mNeonGreen-tagged Tet-inducible strains with Hoechst DNA staining. Cells were imaged after 16 hours of doxycycline treatment in -Leu media and after 8 hours of subsequent aggregation selection in -Leu -Ura after doxycycline removal.

Demonstrating the utility of the system, the well-characterized Sup35 *[PSI+]* prion grew on selective media following doxycycline removal (Figure 2B). However, in the absence of doxycycline, only a few clonal outgrowths were observed. Cells expressing wild-type *IES6* also grew, albeit to a lesser degree than Sup35, perhaps due to the shorter length of the Ies6 PLD, which is 23 amino acids (Figure 1A) compared to that of Sup35, which is 123 amino acids. Removal of Ies6’s PLD in *ies6-Cterm* cells resulted in decreased growth, whereas, conversely, expression of the PLD alone resulted in increased growth compared to wild-type cells. Importantly, cells lacking the Hsp70 chaperones, Ssa1 and Ssa2, exhibited increased growth in selective media, which is particularly predominant in the *ies6-PLD* mutant (Figure 2B). Additionally, brief heat shock (1 hour) of cells to induce global heat shock protein expression, prior to plating on selective media, resulted in decreased growth (Figure 2C). Further emphasizing the relationship between Ies6 and prion-regulating heat shock proteins.

We next investigated subcellular localization of Ies6-mNeonGreen in the Tet-induced aggregation selection system. We found that doxycycline-induced overexpression of both wild-type Ies6 and Ies6-Cterm lacking the PLD, resulted in nuclear localization (Figure 2D). Wild-type Ies6 was found to form nuclear foci, while Ies6-Cterm protein diffusely overlapped with the DNA stain. However, overexpression of Ies6-PLD alone resulted in aggregate formation outside the nucleus. In selective -Leu -Ura media without doxycycline, wild-type Ies6 was observed to be primarily nuclear (Figure 2D, right panels). Expression of PLD alone was largely diffuse throughout the cell, with smaller aggregate formations compared to the previous doxycycline-induced overexpression. Cells expressing Ies6-Cterm alone had reduced growth during selection and were not imaged.

### Arp5 regulates aggregation of Ies6

As previously mentioned, Ies6 is both a subunit in the INO80 complex and part of an abundant independent subcomplex with Arp5 (Yao et al. 2015, 2016). Purification of Arp5 recapitulates previous results of Ies6 association with Arp5 and the INO80 complex, and deletion of Ies6’s PLD does not disrupt this association (Figure 3A). As before, the Ies6-PLD alone is not visible due to its small size, yet its expression disrupts Arp5’s association with the INO80 complex in a manner similar to that of complete *IES6* deletion.

**Figure 3.**
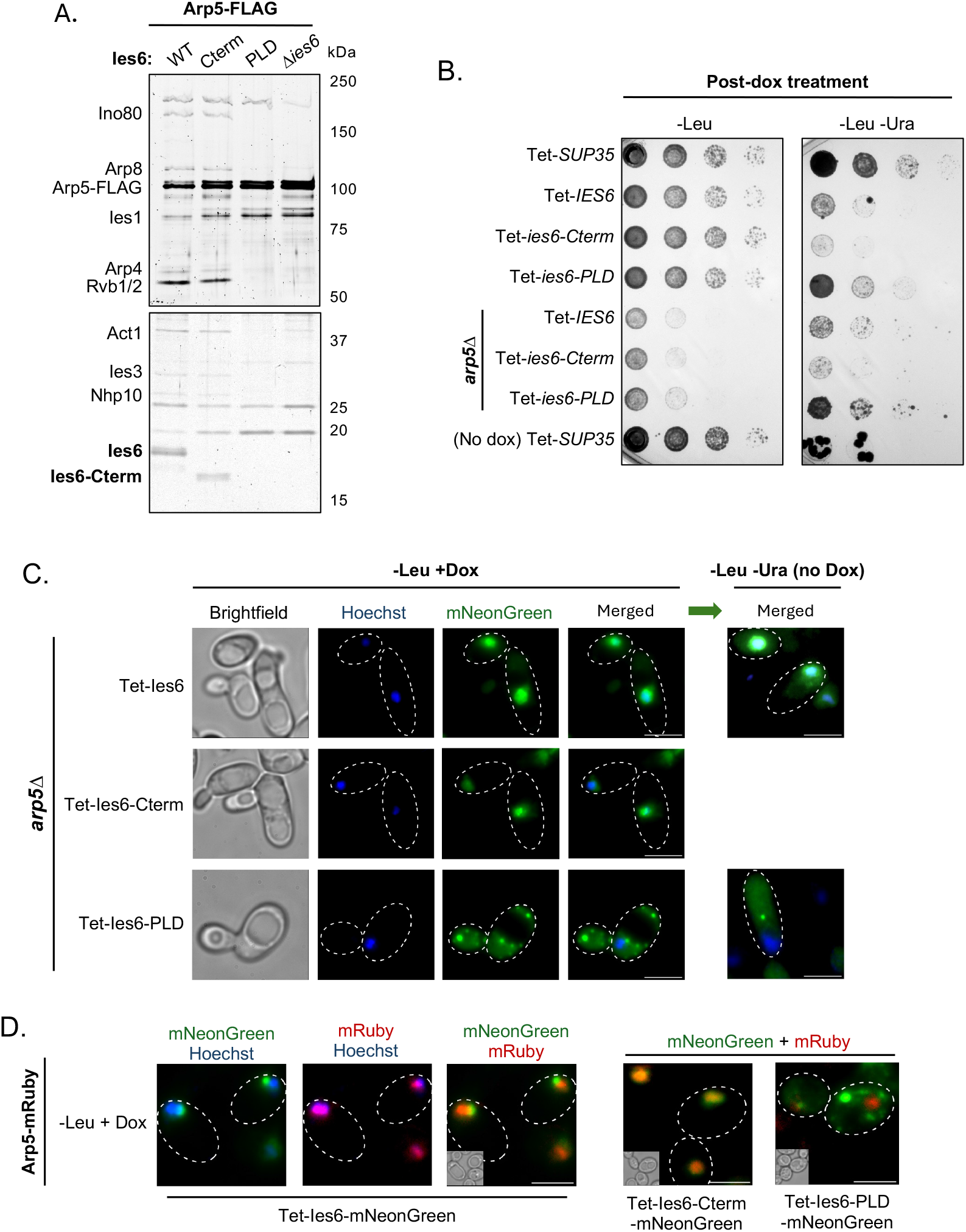
Deletion of *ARP5* increases Ies6 PLD aggregation. **(A)** Arp5-Flag purification with the indicated *ies6* mutants. Top, 6% SDS-PAGE gel and bottom, 13.5% SDS-PAGE gel. Previously identified INO80 subunits (Shen et al. 2000; Yao et al. 2015, 2016) are indicated on the side of the top 6% SDS-PAGE gel and the bottom 13.5% gel. Note that Arp5 forms a subcomplex with Ies6 alone, thus other INO80 subunits are less prominent. **(B)** Growth of wild-type and *arp5Δ* mutants expressing indicated Tet-inducible constructs on both -Leu (no aggregation selection) and -Leu -Ura (aggregation selection). All strains were treated with doxycycline prior to plating, except where indicated for the (no dox) negative Sup35 control. **(C)** Fluorescent imaging of indicated Tet-inducible mNeonGreen constructs in *arp5Δ* strains with Hoechst DNA staining. Cells were imaged after 16 hours of doxycycline treatment in -Leu media and after 8 hours of subsequent aggregation selection in -Leu -Ura after doxycycline removal. Scale bar indicates 5 microns. **(D)** Imaging of Arp5-mRuby and indicated Tet-inducible mNeonGreen constructs. Cells were treated as in (C).

We again utilized the aggregation selection assay to determine the effect of Arp5 on Ies6 PLD-induced aggregation. We confirmed that deletion of *ARP5* substantially decreased fitness on non-selective media (Figure 3B), as previously published (Yao et al. 2015, 2016). However, on selective media after doxycycline removal, we found that the growth of cells expressing Ies6 constructs was similar in both wild-type and *arp5Δ* cells (Figure 3B). Thus, when accounting for the decrease in baseline growth of *arp5Δ* mutants, these cells exhibited relatively more growth than wild-type cells during selection, indicating a greater propensity for Ies6 aggregation in cells lacking *ARP5*.

Fluorescent assays were performed to examine subcellular localization of Ies6-mNeonGreen in *arp5Δ* cells using the Tet-induced aggregation selection system. As in wild-type cells, Ies6 colocalized with the nucleus during doxycycline induction (Figure 3C). Aggregate formation of the PLD alone was again prominent in *arp5Δ* cells during doxycycline induction. Following growth in selective media, while wild-type Ies6 largely colocalized with DNA stain, persistent aggregates outside of the nucleus were observed in cells expressing Ies6-PLD alone (Figure 3C, right panels).

We then examined endogenous Arp5-mRuby localization in the Tet system during doxycycline induction and found that Arp5-mRuby colocalized with Hoechst DNA stain and appeared distinct, yet proximal, to wild-type Ies6 foci formation (Figure 3D). As before, overexpressed Ies6-PLD formed cytosolic aggregates, which did not colocalize with the nuclear Arp5 (Figure 3D, right panel). However, in Ies6-Cterm cells, the Ies6 mutant lacking the PLD strongly colocalizes with nuclear Arp5. Collectively, these results indicate that Arp5 does not overlap with Ies6 aggregate formation and, furthermore, *ARP5* deletion promotes Ies6 aggregation. Thus, suggesting that Arp5 association with Ies6 may restrain Ies6 PLD-mediated aggregation.

### Ies6 aggregation modifies subtelomeric and metabolic gene expression

In order to determine the impact of PLD-mediated Ies6 aggregation on gene expression, RNA-seq was performed on cells expressing wild-type *IES6* and the *ies6-Cterm* mutant constructs in the Tet aggregation system. Following doxycycline induction and removal, cells were grown in - Leu -Ura media to select for Ies6 aggregation. Selection was performed on solid media rather than in liquid to prevent rapid outgrowth of colonies that circumvent aggregate selection. Although the Ies6-Cterm expressing cells do not grow well in -Leu -Ura media, they served as a background control for cells undergoing selection. When comparing *ies6-Cterm* expressing cells lacking the PLD to wild-type *IES6* during aggregate selection, we found over 3,000 genes that were significantly differentially expressed, including both up- and down-regulated genes (Figure 4A).

**Figure 4.**
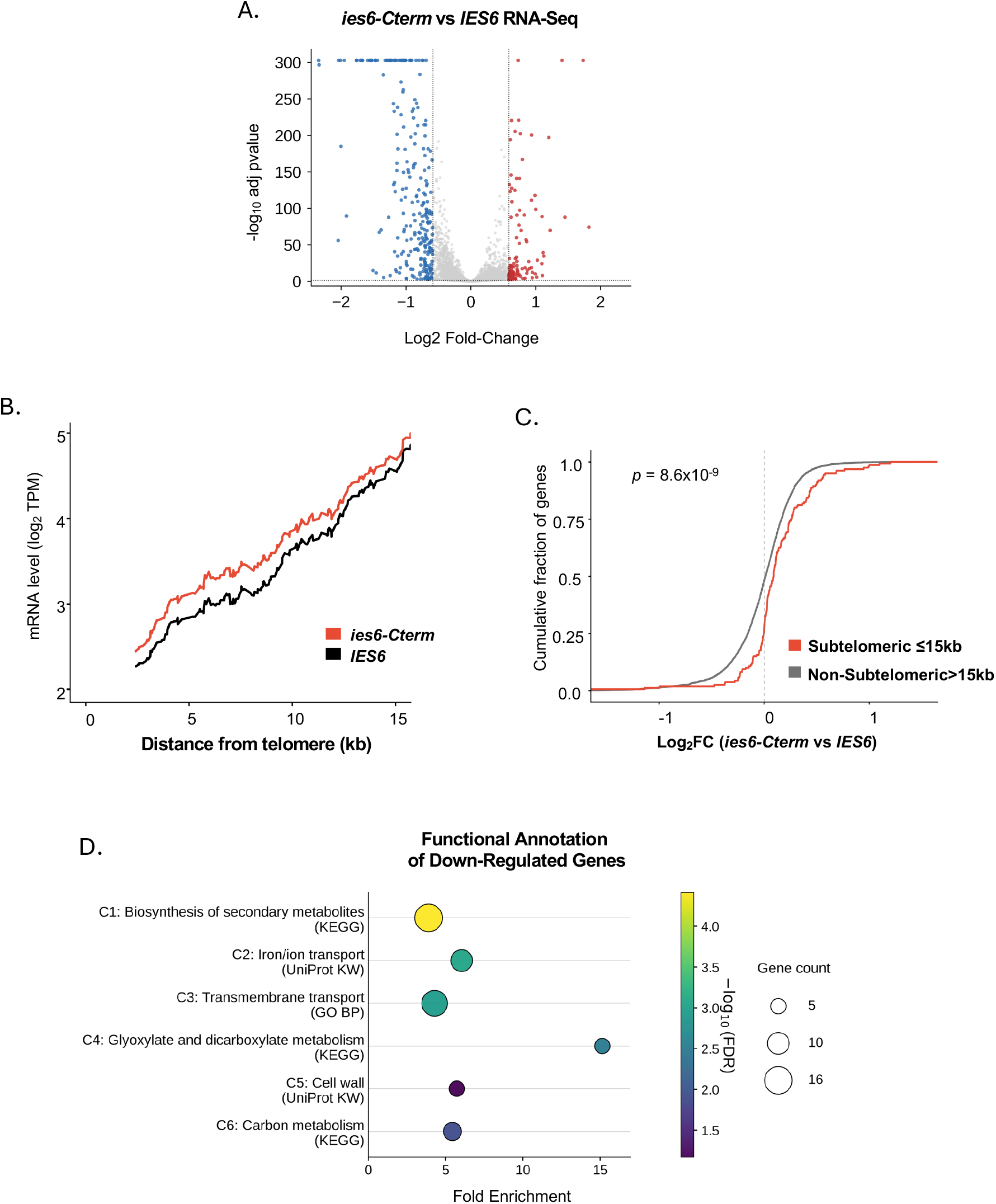
Loss of Ies6 aggregation results in metabolic gene expression changes. RNA-seq comparing Tet-inducible *ies6-Cterm* vs wild-type (WT) *IES6* strains after selection of Ies6 aggregation following doxycycline removal and growth on -Leu -Ura plates. **(A)** Volcano plot of gene expression changes. **(B)** The rolling average of log2(TPM + 1) plotted as a function of the distance from 0-15 kb from the telomere (window = 100 genes, step-size = 1 gene). **(C)** The empirical cumulative distribution function (ECDF) of log2 fold-changes for subtelomeric genes (≤15 kb from telomere; N = 160) and non-subtelomeric gene distribution (>15 kb; N = 5,810) across log2 fold change values (two-sided Kolmogorov–Smirnov test: D = 0.249, *p* = 8.6x10^-9^).**(D)**DAVID functional annotation enrichment of significantly downregulated genes with log2 fold change greater than -/+1. Representative terms from each cluster (C1-C6) are shown with the corresponding database source.

Prior work has shown that the INO80 complex maintains the boundary between euchromatin and repressed subtelomeric regions (Xue et al. 2015). To examine whether disruption of the PLD affects subtelomeric gene expression, we compared the subtelomeric (0-15kb) RNA-seq profiles of *ies6-Cterm* mutant and wild-type *IES6* across all chromosomes. Interestingly, loss of Ies6’s PLD resulted in an increase in subtelomeric gene expression relative to wild-type Ies6 (Figure 4B). Likewise, the fold-change differences between *ies6-Cterm* and *IES6* expressing cells were significantly higher in subtelomeric regions compared to other genomic regions, indicating that loss of Ies6’s PLD results in reduced subtelomeric repression (Figure 4C).

When examining the functional annotation of all significantly differentially expressed genes, no significant enrichment was observed among up-regulated (L2FC>1) genes. However, down-regulated (L2FC<1) genes were enriched in metabolite regulation, glucose starvation, and respiratory transcriptional programs (Figure 4D). These trends are particularly notable, given that the cells were grown in glucose-replete conditions, and suggest that aggregation of Ies6 may prime transcriptional pathways involved in adaptive carbon stress responses.

## DISCUSSION

In this study, we report that the Ies6 subunit of the INO80 complex contains a large domain of predicted disorder that surrounds a smaller N-terminal prion-like domain (PLD). This domain promotes aggregation in fluorescent assays and a growth selection system that enriches for protease-resistant assemblies. While deletion of the PLD reduces aggregation and alters carbon metabolism pathways. Collectively, the results of this study identify Ies6 as a prion-like INO80 subunit that undergoes conformational changes to regulate chromatin-regulated transcription of metabolic genes.

The identification of prion-like behavior in Ies6 is notable given the specialized role of the Arp5– Ies6 in chromatin remodeling function. Our previous work demonstrated that Arp5-Ies6 is both a module of the INO80 complex and an abundant subcomplex in yeast (Yao et al. 2015, 2016). Notably, using *in vitro* assays, the purified Arp5-Ies6 subcomplex can assemble into the INO80 complex to stimulate chromatin-remodeling activities. Thus, Ies6 aggregation may modify the availability of Arp5-Ies6 to assemble with the INO80 complex. In addition, it may alter INO80 chromatin remodeling function if aggregation is stimulated while associated with the INO80 complex. For example, Ies6 aggregation may impact INO80 substrate specificity. INO80 has been shown to remodel hexasomes preferentially over canonical nucleosomes (Hsieh et al. 2022; Zhang et al. 2023), suggesting that INO80 may be particularly important at dynamic chromatin regions where H2A–H2B dimers are exchanged or partially displaced. Because Arp5–Ies6 stimulates core remodeling activity, prion-like regulation of Ies6 could alter INO80 substrate specificity and function at chromatin loci enriched for hexasomes. Future assays investigating Ies6 and INO80 colocalization at hexasomes may shed light on these possibilities.

Importantly, conditions that naturally induce Ies6 aggregation are still under investigation. However, it is notable that loss of Ies6 aggregation results in decreased expression of gene pathways associated with nonfermentable carbon utilization and respiratory metabolism, even in the presence of ample glucose. This is somewhat similar to the [*GAR+*] prion-induced state, which relaxes glucose repression, allowing yeast to use different carbon sources even while in the presence of glucose (Brown and Lindquist 2009; Jarosz et al. 2014; Garcia et al. 2016). Thus, it may be that Ies6 undergoes aggregation to allow metabolic flexibility, thereby promoting survival in fluctuating nutrient environments.

Other chromatin-associated prions provide additional relevant comparisons. Swi1, a subunit of the SWI/SNF chromatin remodeling complex, forms the [*SWI+*] prion, which alters SWI/SNF-dependent gene regulation and affects growth on non-glucose carbon sources (Du et al. 2020, 2008). Similarly, Snt1, the scaffold subunit of the Set3C histone deacetylase complex, forms the [*ESI+*] prion, which establishes transgenerational inheritance of active subtelomeric chromatin state (Harvey et al. 2020). As in these examples, Ies6 may provide a mechanism by which a chromatin regulator adopts an alternative protein state to alter gene expression. However, a notable difference from these other prions is that, while the PLD of Ies6 promotes aggregation, non-Mendelian inheritance has not yet been directly examined.

Nevertheless, the results shown in this manuscript demonstrate that the prion-like properties of Ies6 regulate gene expression at multiple distinct genomic locations. Specifically, removal of the prion-like domain results in the de-repression of subtelomeric genes, consistent with prior work on INO80 in regulating heterochromatin-euchromatin boundaries (Xue et al. 2015). In addition, the prion-like domain of Ies6 regulates metabolic gene expression, which ultimately affects adaptive stress responses. As previously mentioned, the INO80 complex has been linked to metabolic homeostasis through TORC1-responsive chromatin regulation, histone acetylation at metabolic genes, mitochondrial function, and the coordination of metabolic gene expression with the nutrient environment (Yao et al. 2016; Gowans et al. 2018; Beckwith et al. 2018). When taken together, these results highlight the Ies6 subunit of the INO80 complex in maintaining homeostasis during environmental adaptation.

## METHODS

### Strain and plasmid generation

All strains are from the *S*.*cerevisiae* S288C BY4741 background using standard techniques for endogenous gene deletion and tagging (Kaiser et al. 1994). The tetracycline overexpression system used the previously developed strong semisynthetic TetO promoter, which is repressed by TetR and controlled by an autorepression loop to limit leaky expression and cell-to-cell variation (Azizoglu et al. 2021). The Tet-inducible plasmids encoded the *LEU2* selection marker on the backbone. The aggregation selection plasmid consisted of a three-part fusion construct with the *URA3* gene, *IES6/SUP35*, followed by the synthetic *E*.*coli* dihydrofolate reductase R12Y/Y100I destabilizing domain, previously described (Iwamoto et al. 2010).

### Fitness and aggregation selection assays

All strains were streaked onto plates overnight prior to growth in fitness and aggregation selection assays. Wildtype and endogenous mutants were inoculated into YPD liquid media overnight, then diluted to OD_660_ of 0.1 prior to serial dilution on YPD plates and growth at 30°C. For cells expressing the Tet-inducible system, cells were grown overnight in synthetic complete -Leu media, followed by inoculation into fresh -Leu media with 1000ng/mL doxycycline for 16 hours. Cultures were then adjusted to OD_660_ of 1.0 and serially diluted 1:6 before spotting on -Leu and -Leu -Ura plates, without doxycycline, and grown at 30°C for 2-3 days. Where indicated, cells were heat shocked at 42°C for 1 hour prior to plating.

### Protein purification

Protein complexes were purified using anti-FLAG affinity beads, as previously described (Shen 2003; Yao et al. 2015, 2016). Briefly, cells were frozen in liquid nitrogen then broken using a commercial blender. Cell lysates were resuspended in HEGN buffer (25 mM HEPES-KOH pH 7.6, 1 mM EDTA, 10% glycerol, 0.02% Nonidet P-40, 2.5 mM DTT, 2 mM MgCl2, protease inhibitors) with 0.5 M KCl. Soluble lysate was obtained by ultracentrifugation and incubated with FLAG affinity-agarose (Sigma). Beads were washed with HEGN buffer and eluted with FLAG peptide. Purified protein was electrophoresed on 6% and 13.5% SDS-PAGE gels, then stained with Oriole Fluorescent Protein Stain (Bio-Rad) before imaging.

### Microscopy

Endogenously tagged strains were inoculated into YPD liquid media overnight, then diluted and grown to mid-log phase before harvesting. For cells expressing the Tet-inducible system, cells were grown overnight in synthetic complete -Leu media, followed by inoculation into fresh -Leu media with 1000ng/mL doxycycline for 16 hours, then diluted to 0.2 OD_660_ in -Leu -Ura liquid media for 8 hours without doxycycline before harvesting.

For Hoechst staining, 1×10^7^ cells in media were incubated with a final concentration of 4% paraformaldehyde while rotating at room temperature for 10 minutes, followed by quenching with 100 mM glycine for 5 minutes. Cells were washed twice with PBS and permeabilized with 0.1% Triton X-100 in PBS for 5 min. Cells were washed again with PBS and resuspended in 1ml PBS with 2ug/mL Hoechst stain while rotating for 15min at room temperature. Cells were washed twice with PBS and imaged using Leica DMi8 fluorescence microscope with a 100x objective.

### RNA-Seq

Tet constructs with wildtype *IES6* or *ies6-Cterm* mutants were transformed into strains with matching wildtype *IES6 or ies6-Cterm* endogenous mutants. Cells were grown overnight in synthetic complete -Leu media. Cells were then split into two groups and both inoculated into fresh -Leu liquid media, with one group receiving 1000ng/mL doxycycline for 16 hours and the other group untreated. Subsequently, 3×10^6^ cells were plated, onto -Leu -Ura for the doxycycline-treated cells and -Leu for the untreated cells and allowed to grow at 30°C for 3 days. Cells were harvested, and an OD_660_ of 1.0 was used to prepare the total RNA using MasterPure Yeast RNA Purification Kit (Biosearch Technologies). RNA quality (RIN > 7) was assessed using Agilent Tape Station, and SIRV-Set 4 spike-in was added according to manufacturer specifications. RNA was prepped using NEBNext Ultra II Directional RNA Library Prep Kit for Illumina with Dual Index UMI Adaptors and sequenced on Illumina NovaSeq X Plus. Reads were processed using nfcore/rna-seq pipeline (Ewels et al. 2020) followed by DESeq2 (Love et al. 2014).

## ACKNOWLEDGMENTS

We thank members of the Morrison and Jarosz labs for advice and reagents. We thank Asli Azizoglu and Roger Bent, PhD, for assistance with the Tet-inducible system. We thank Sean Beckwith, PhD, for helpful discussions during the initial stages of this research.

## AUTHOR CONTRIBUTIONS

Conceptualization - AJM; methodology - DCE, KCG, DFJ, and AJM; investigation – DCE, KCG and AJM; writing original draft - AJM; writing, review, and editing – DCE, KCG, and AJM; funding acquisition - AJM; resources - AJM; supervision - AJM.

